# Chromosome-level genome assembly for the angiosperm *Silene conica*

**DOI:** 10.1101/2023.09.05.556365

**Authors:** Peter D. Fields, Melody M. Weber, Gus Waneka, Amanda K. Broz, Daniel B. Sloan

## Abstract

The angiosperm genus *Silene* has been the subject of extensive study in the field of ecology and evolution, but the availability of high-quality reference genome sequences has been limited for this group. Here, we report a chromosome-level assembly for the genome of *Silene conica* based on PacBio HiFi, Hi-C and Bionano technologies. The assembly produced 10 scaffolds (one per chromosome) with a total length of 862 Mb and only ∼1% gap content. These results confirm previous observations that *S. conica* and its relatives have a reduced base chromosome number relative to the genus’s ancestral state of 12. *Silene conica* has an exceptionally large mitochondrial genome (>11 Mb), predominantly consisting of sequence of unknown origins. Analysis of shared sequence content suggests that it is unlikely that transfer of nuclear DNA is the primary driver of this mitochondrial genome expansion. More generally, this assembly should provide a valuable resource for future genomic studies in *Silene*, including comparative analyses with related species that recently evolved sex chromosomes.

**Significance:** Whole-genome sequences have been largely lacking for species in the genus *Silene* even though these flowering plants have been used for studying ecology, evolution, and genetics for over a century. Here, we address this gap by providing a high-quality nuclear genome assembly for *S. conica*, a species known to have greatly accelerated rates of sequence and structural divergence in its mitochondrial and plastid genomes. This resource will be valuable in understanding the coevolutionary interactions between nuclear and cytoplasmic genomes and in comparative analyses across this highly diverse genus.

## Introduction

*Silene* (Caryophyllaceae) is a diverse angiosperm genus that encompasses over 800 species and has been the subject of extensive study in ecology and evolutionary genetics (Bernasconi, et al. 2009; Jafari, et al. 2020). Species in this genus have served as models for investigating topics such as breeding system and sex chromosome evolution (Nicolas, et al. 2005; Kejnovsky and Vyskot 2010; Muyle, et al. 2012; Krasovec, et al. 2018; Martin, et al. 2019), host-pathogen interactions (Alexander and Antonovics 1988; Kephart, et al. 2006; Le Gac, et al. 2007), heavy metal tolerance (Antonovics, et al. 1971; Bringezu, et al. 1999; Papadopulos, et al. 2021), species invasion (Blair and Wolfe 2004; Keller and Taylor 2008; Keller, et al. 2014), and cytoplasmic/cytonuclear genetics (Taylor, et al. 2001; Touzet and Delph 2009; Sloan, Alverson, Chuckalovcak, et al. 2012; Fields, et al. 2014; Havird, et al. 2015). Even though many of these topics are directly rooted in areas of genome biology, the availability of high-quality genomic resources for *Silene* has remained limited. An extensive number of RNA-seq projects and transcriptome assemblies have been performed in *Silene* (e.g., Blavet, et al. 2011; Havird, et al. 2017; Jafari, et al. 2020; Warren, et al. 2022), but initial whole-genome sequencing efforts in *Silene* were limited to a mix of short-read and early-generation long-read technologies, resulting in highly fragmented and incomplete assemblies (Papadopulos, et al. 2015; Krasovec, et al. 2018; Williams, et al. 2021). Only recently was the first chromosome-level genome assembly produced for a member of this genus (Yue, et al. 2023).

To date, genome assemblies have been published for only two *Silene* species: *S. latifolia* (Papadopulos, et al. 2015; Krasovec, et al. 2018; Yue, et al. 2023) and *S. noctiflora* (Williams, et al. 2021). These two species are found near the high end of the known range of 0.7-3.3 Gb for genome sizes of diploid species in this genus (Pellicer and Leitch 2019). At the other end of the spectrum, *S. conica* has one of the smallest estimated genome sizes (0.9 Gb) in *Silene* (Williams, et al. 2021).

*Silene conica* and its relatives within section *Conoimorpha* also have a reduced chromosome number (2n=20) compared to the ancestral state (2n=24) for *Silene* (Bari 1973). In contrast to its small nuclear genome size, *S. conica* has one of the largest mitochondrial genomes of any eukaryote at >11 Mb, containing over 99% non-coding content of largely unknown origin (Sloan, Alverson, Chuckalovcak, et al. 2012). The cytoplasmic genomes in *S. conica* also exhibit other distinctive features, including accelerated evolutionary rates, major structural changes, and extensive gene loss (Erixon and Oxelman 2008; Sloan, Alverson, Chuckalovcak, et al. 2012; Sloan, Alverson, Wu, et al. 2012). As such, this species has been a valuable model for studying how changes in cytoplasmic genomes can spur cytonuclear coevolution (Rockenbach, et al. 2016; Havird, et al. 2017; Abdel-Ghany, et al. 2022).

With the continuing improvement of DNA sequencing technologies, it is becoming increasingly possible to generate chromosome-level assemblies, even for complex eukaryotic nuclear genomes like those of plants (Jiao and Schneeberger 2017; Belser, et al. 2018; Shirasawa, et al. 2021). In particular, the advent of Pacific Bioscience (PacBio) HiFi sequencing has resulted in a major step forward, providing single-molecule long reads (∼15-25 kb) at high accuracy (>99%) (Wenger, et al. 2019). Here, we report a chromosome-level assembly of the *S. conica* genome generated from HiFi sequencing in combination with Bionano optical mapping (Lam, et al. 2012) and Hi-C scaffolding (Belton, et al. 2012).

## Results & Discussion

### Chromosome-Level Genome Assembly

PacBio HiFi sequencing of *S. conica* total-cellular DNA produced 2.23M circular consensus sequence (CCS) reads with an average length of 14.9 kb and a total yield 33.18 Gb. Our base assembly of these reads produced by *hifiasm* was a total of 938 Mb in length, with an N50 length of 14.95 Mb (n=20), a maximum contig length of 46.80 Mb, and a total of 1367 contigs. Following the application of purge_haplotigs, we saw improvement in assembly metrics, including an N50 length of 16.23 Mb (n=18), a total contig number of 217, and a total assembly length of 869 Mb, which is slightly shorter than the previous estimate of 930 Mb from flow cytometry (Williams, et al. 2021). This base assembly was then used as an input for Bionano Access.

A single round of hybrid scaffolding with a Bionano optical map resulted in a large improvement in overall genome contiguity. Specifically, while the overall genome did not change substantially in length (862 Mb), Bionano scaffolding produced 16 scaffolds with an N50 scaffold length of 74.70 Mb (n=5), a longest scaffold of 122.47 Mb, and 65 gaps totaling only 6.42 Mb. This small amount of gapped content (∼1% of the assembly) compares favorably to the level of contiguity achieved for most chromosome-level assemblies of plant genomes (Shirasawa, et al. 2021). We then proceeded with analysis of Hi-C data to generate chromosomal scaffolds, although six of the scaffolds already appeared to represent near or whole chromosomes at this stage.

Visualization of the Hi-C based contact network (Figure S1) provided support for joining the above Bionano scaffolds into the expected number of 10 chromosomes (Bari 1973). However, this visualization also revealed that the Bionano scaffold containing Chr1 was misassembled and contained a large portion of Chr6 (Figure S1). The point at which these two regions were joined corresponded to a large gap in the Bionano scaffold that was associated with highly repetitive ribosomal DNA. After breaking this misassembly and joining the Bionano scaffolds based on the Hi-C contract map, our analysis resulted in 10 chromosome-level scaffolds with 71 gaps, a total length of 862 Mb, and 37.0% GC content (Figure S2).

### Genome Annotation

We used BUSCO (Manni, et al. 2021) to assess the biological completeness of our genome assembly. At present, there is no Caryophyllaceae-specific ancestral gene set, so we used the embryophyta_odb10 dataset for the BUSCO analysis, which produced a completeness score of 97.6. We detected 1586 of the 1614 BUSCO genes searched (1506 complete and single-copy, 70 complete but duplicated, and 10 fragmented), while 28 BUSCOs were missing. An ancestral gene set derived from species that better represent genomes of the Caryophyllaceae might result in a slightly higher overall BUSCO score.

In order to maximize our genome annotation completeness, we relied on a plurality of approaches. Specifically, we used a combination of MAKER2 (Holt and Yandell 2011), Funannotate (Palmer and Stajich 2020), and BRAKER (Hoff et al. 2019) individually, followed by the merging and collapsing of redundant annotations using AGAT (Dainat et al. 2023). For MAKER2, we utilized an iterative application of the pipeline to annotate the *S. conica* genome. Following the first round of MAKER2, which relied on protein and transcript hints alone, we identified a total of 51,311 putative gene models. Our second round of MAKER2, which included the application of AUGUSTUS and SNAP *ab initio* hints, as well as the gene models from MAKER2 round one, resulted in 57,309 putative gene models. Our third round of MAKER2, which incorporated GeneMark *ab initio* hints, as well as the gene models from MAKER2 round two, resulted in 56,305 putative gene models. Finally, the fourth round of MAKER2 included the application of AUGUSTUS and SNAP *ab initio* hints, which were trained off gene models resulting from MAKER2 round three, and also included gene models from MAKER2 round three. This final round resulted in 58,409 putative gene models, which were filtered based upon an annotation edit distance (AED) score threshold of ≤ 1, yielding a total of 47,262 putative gene models. Our Funannotate annotation resulted in a total of 55,798 putative gene models. Finally, BRAKER annotation resulted in a total of 56,992 putative gene models. Our merging and de-duplication of gene models using AGAT resulted in a total of 63,211 putative gene models. The resultant BUSCO score of our annotation is nearly as good as the overall genome with a completeness score of 93.2%, suggesting our automated annotation process was highly effective. Manual curation would likely improve the overall accuracy of individual gene models, and filter out spurious annotations that may have inflated the number of identified gene models.

Karyotyping has indicated that the *S. conica* chromosomes are metacentric or submetacentric (Bari 1973). Accordingly, the chromosomes show the typical pattern of higher gene densities and lower CpG methylation rates at the ends of chromosome arms relative to the middle of the chromosome (Figure 1). However, it should be noted that this expectation was used to orient scaffolds during the final Hi-C based joining for Chr3 and Chr8 (see Methods). Therefore, the gene density patterns for these two chromosomes do not provide any further independent evidence for a typical metacentric structure.

**Figure 1.**
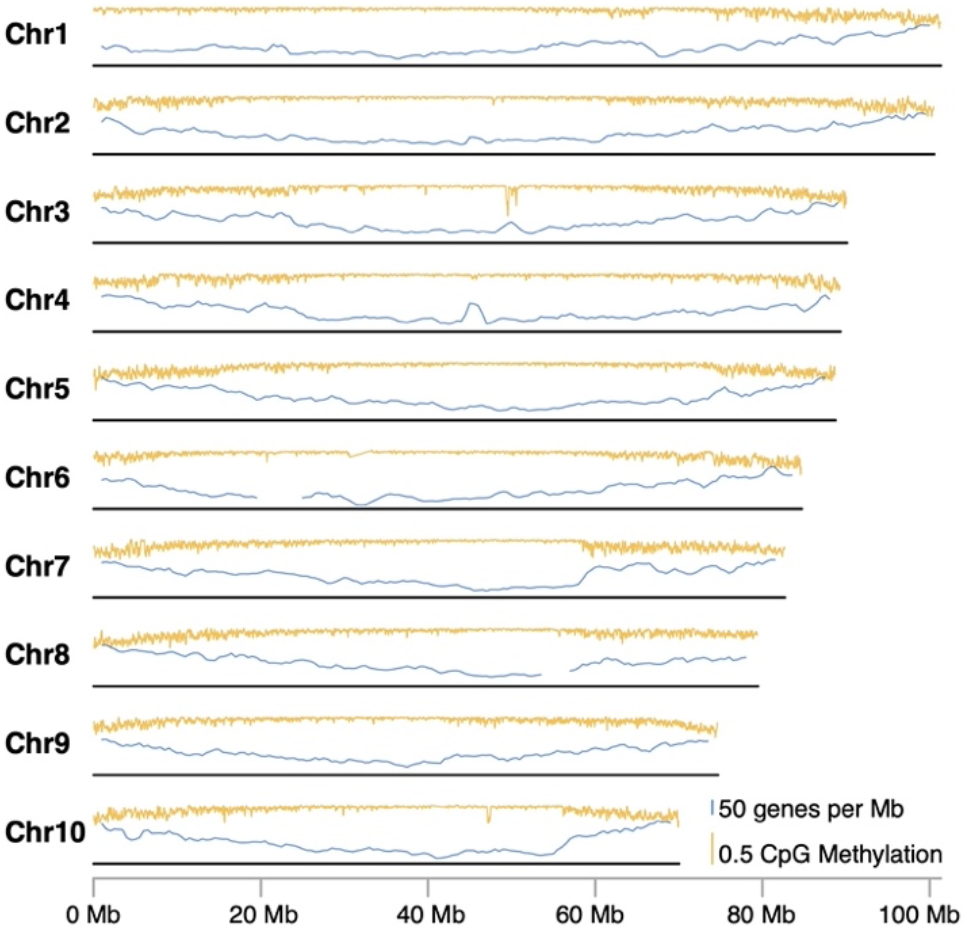
Summary of CpG methylation rates and gene density across the length of each chromosome in the *S. conica* genome. The orange trace represents a sliding window summary (100-kb window size and 25-kb step size) of the proportion of CpG sites inferred to be methylated from analysis of PacBio read data. The blue trace represents a sliding window summary (2-Mb window size and 500-kb step size) of the average number of annotated genes per Mb. The localized dips in estimated methylation rates on Chr3 and Chr10 correspond to the positions of large *numt* insertions (Figure 3). CpG methylation generally does not occur in mitochondrial DNA, so these dips are likely an artifact result from incorrectly mapping true mitochondrial reads to these *numt* regions (Fields, et al. 2022). Gene density estimates are not reported for the regions on Chr6 and Chr8 with tandemly repeated ribosomal DNA.

An analysis of interspersed repeat content in the genome found that transposable elements constituted approximately three-quarters of all sequence content (Table S1). Indeed, long terminal repeat (LTR) retrotransposons by themselves were estimated to account for more than half of the genome.

### DNA transfer between nuclear and cytoplasmic genomes

The insertion of mitochondrial and plastid DNA into the nuclear genome (known as *numts* and *nupts*, respectively) is a widespread phenomenon across eukaryotes (Hazkani-Covo, et al. 2010; Zhang, et al. 2020). In plants, the movement of DNA in the opposite direction – from the nucleus to cytoplasmic genomes – is also common for mitochondria (Goremykin, et al. 2012; Qiu, et al. 2014) but very rare for plastids (Smith 2014). To characterize the extent of intracellular DNA transfer in *S. conica*, we used BLAST searches to compare the mitochondrial and plastid genomes against our nuclear genome assembly. We found regions with significant similarity to the cytoplasmic genome were widely distributed across the nuclear genome (Figure 2). A total of 1938 kb of nuclear DNA sequence (0.22% of the nuclear genome) shared similarity with the mitochondrial genome, and 186 kb shared similarity with the plastid genome (0.02% of the entire nuclear genome). This amount of shared content is well within the range observed in other species. Angiosperm nuclear genomes have been estimated to share anywhere from 142 kb to 11,420 kb of sequence with the mitochondrial genome (covering between 0.03% and 2.08% of the assembled nuclear genomes) and 36 kb to 9830 kb with the plastid genome (0.01% to 1.49% of the assembled nuclear genomes) (Zhang, et al. 2020).

**Figure 2.**
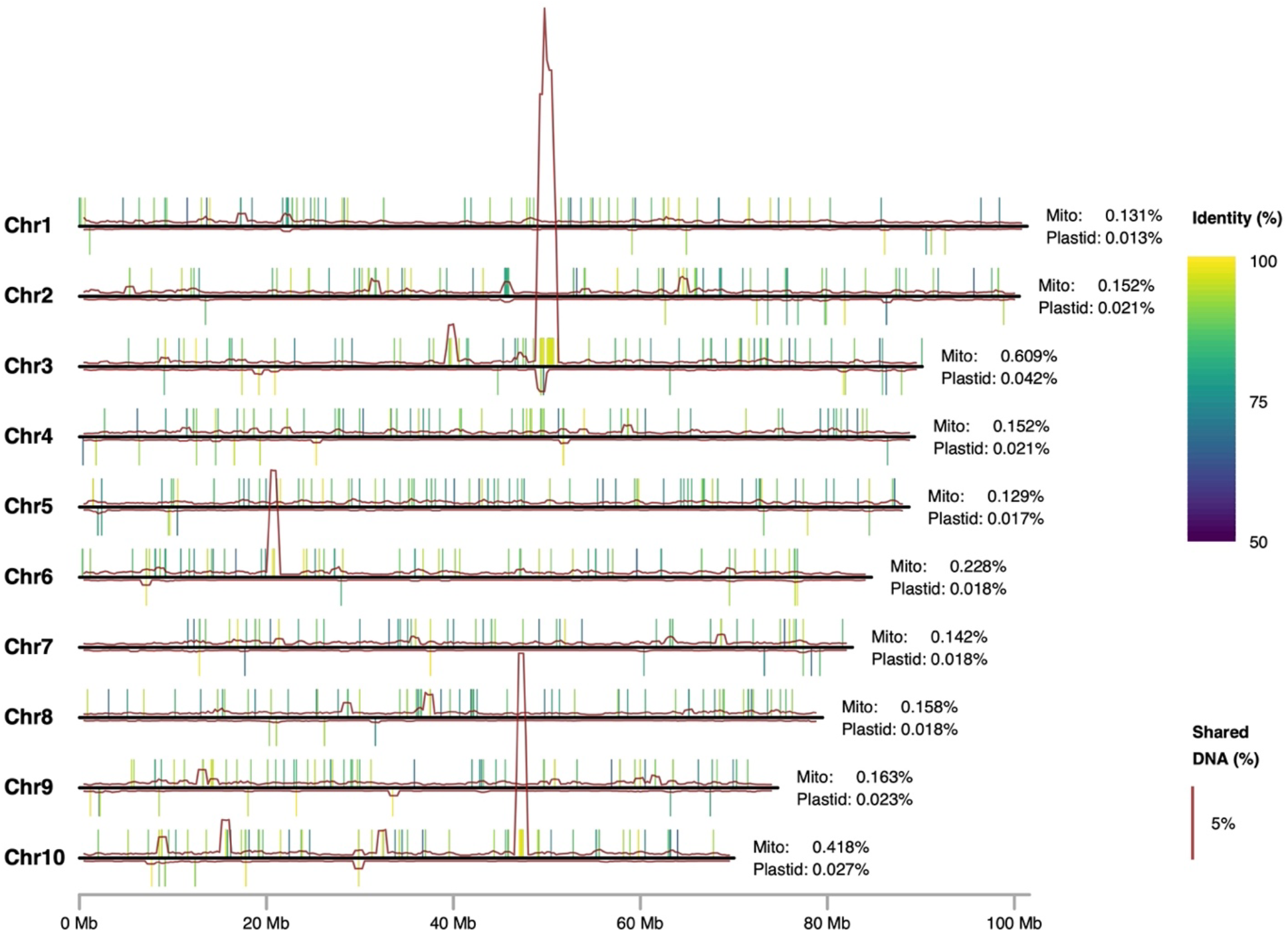
Summary of sequence content shared between the *S. conica* nuclear and cytoplasmic genomes. Tick marks above and below each nuclear chromosome indicate sequence content shared with the mitochondrial and plastid genomes, respectively, as identified by BLAST analysis (minimum hit length of 300 bp and e-value threshold of 1e-6). The color of each tick mark indicates the percent nucleotide identity of the BLAST hit. The red traces are from a sliding window analysis (1-Mb window size and 250-kb step size), indicating the percentage of sequence in the corresponding window that is shared with the mitochondrial genome (above the chromosome) or plastid genome (below the chromosome). Values on the right of each chromosome indicate the overall percentage of sequence shared with each cytoplasmic genome.

The shared sequences in *S. conica* represented a total of 2863 kb (25.3%) in the mitochondrial genome and 103 kb (70.3%) in the plastid genome. Note that the raw totals represented by these shared sequences differ between the nuclear and cytoplasmic genomes because of the differences in the extent to which the shared content is repeated within each of the genomes. There were two regions of the nuclear genome that were especially rich in DNA shared with the mitochondrial genome (Figure 2): the ∼1405 kb region Chr3 from position 49.307 Mb to 50.712 Mb and the ∼335 kb region on Chr10 from position 47.115 Mb to 47.450 Mb. Although synteny in these regions of shared sequence was highly fragmented relative to the mitochondrial genome, much of the sequence retained very high nucleotide identity (>99% in many cases) between the two genomes, implying that the transfer(s) responsible for the shared content occurred relatively recently.

Given the rarity with which foreign DNA is inserted into plastid genomes (Smith 2014), it is reasonable to interpret any regions of shared sequence between nuclear and plastid genomes as being ultimately plastid in origin (i.e. *nupts*). However, because transfer between mitochondrial and nuclear genomes is bidirectional in plants, it is more challenging to polarize the movement of DNA shared between these two genomes. The fact that approximately one-quarter of the mitochondrial genome content in *S. conica* is shared with the nucleus could mean that nuclear DNA is a major contributor to mitochondrial genome expansion in this lineage (Sloan, Alverson, Chuckalovcak, et al. 2012), as suggested in other plant mitochondrial genomes (Goremykin, et al. 2012). However, it is possible or even likely that most of this shared content results from transfers in the opposite direction given the enormous quantity of *numts* often found in plant nuclear genomes (Zhang, et al. 2020; Fields, et al. 2022). In principle, the direction of transfer could be polarized by identifying homologous content in either the nuclear or mitochondrial genomes of other plant species. However, this is not currently feasible because only a tiny fraction of the *S. conica* mitochondrial genome has detectable homology with any known sequence (Sloan, Alverson, Chuckalovcak, et al. 2012). As such, deciphering the origins of these shared sequences will likely require additional genome sequences from close relatives within *Silene* section *Conoimorpha* or population genomic sampling within *S. conica*.

## Materials and Methods

### Plant material and growth conditions

Seeds from the *S. conica* ABR (Abruzzo, Italy) line were sown in ProMix BX soil mix. The seeds were collected from a sibling of the plant previously used for Iso-Seq analysis (Warren, et al. 2022), and they were the same batch used for ddPCR validation of genome copy number in Broz, et al. (2021). This line had gone through four generations of selfing in the lab and likely started with low levels of heterozygosity given our observation that *S. conica* readily self-fertilizes without any intervention. They were germinated in a growth room under 10-hr short-day lighting conditions and switched to 16-hr long-day conditions after 3 weeks of growth. Lighting was provided with Fluence LEDs at ∼100 µE m^-1^ s^-1^. Plants were initially each grown in a 2-inch pot and then transplanted to 4-inch pots after 4 months of growth. They were watered on an as-needed basis and treated with dilute Miracle-Gro after 5 weeks of growth. Spinosad and neem oil were applied to limit observed outbreaks of thrips and other potential plant pests. After 5 months of growth, 2.3 g of tissue from young rosette leaves was harvested from a single individual and flash frozen in liquid N_2_ for subsequent DNA extraction and PacBio HiFi sequencing. The same plant was allowed to regrow leaf tissue for another 1.5 months, at which point 3.1 g of young rosette leaf tissue and developing shoots (the plant had begun to bolt) were harvested and flash frozen in liquid N_2_ for subsequent Bionano optical mapping. Finally, after two additional weeks of regrowth, 0.6 g of young leaf tissue was harvested from the same plant and immediately used to perform Hi-C library preparation.

### PacBio HiFi sequencing

Flash-frozen leaf tissue (see above) was shipped on dry ice to the Arizona Genomics Institute at the University of Arizona. DNA was extracted with a modified CTAB protocol (Doyle and Doyle 1987). Extracted DNA was analyzed with pulsed-field gel electrophoresis to confirm that it was high molecular weight. A Covaris G-Tube was then used to shear 10 µg of DNA to a size range of ∼10-30 kb followed by bead purification with PB Beads (PacBio). The HiFi sequencing library was constructed following manufacturer’s protocols using SMRTbell Express Template Prep Kit 2.0. The final library was size selected with a range of 10-25 kb on a Blue Pippin (Sage Science) using S1 marker. The recovered final library was quantified with a Qubit HS dsDNA kit, and size was confirmed with an Agilent Femto Pulse system. Sequencing was performed on PacBio Sequel II, using standard manufacturer’s protocols for the Sequel II Sequencing Kit 2.0. The library was sequenced on 2 SMRT Cells (8M) in CCS mode for 30 hours. Analysis was performed with SMRT Link v10.1 software, requiring a minimum of 3 passes for CCS generation.

### Hi-C library preparation and sequencing

Fresh leaf tissue (see above) was used to generate a Hi-C library with the Proximo Hi-C Plant Kit from Phase Genomics (v4.0 protocol). Input material was ground in a mortar and pestle under liquid N_2_. The library was amplified with 12 cycles of PCR, and a total of 234M read pairs (2×150 bp) were generated on an Illumina NovaSeq 6000 platform at the Genomics and Microarray Core at the University of Colorado Anschutz Medical Campus.

### Bionano optical mapping

Flash-frozen leaf tissue (see above) was shipped on dry ice to the McDonnell Genome Institute at Washington University in St. Louis. DNA was isolated with the Bionano plant tissue hybrid protocol (liquid N_2_ grinding and tissue ruptor), including density gradient purification of nuclei, which were then embedded in agarose plugs prior to DNA extraction. Labeling was performed with a Bionano DLS Kit followed by analysis on a Bionano Saphyr platform, generating an estimated genome coverage of 214×. Computational analysis was performed with Bionano Access software.

### Hifiasm de novo assembly

We used the *hifiasm* v.0.15.2-r334 (Cheng, et al. 2021) assembler to generate contigs from PacBio HiFi sequencing data. Given that the focal genotype was relatively inbred, we included the ‘-l0’ flag as part of the assembler configuration, thereby disabling automatic duplication purging. Additional purging of contigs that result from individual heterozygosity, so-called ‘haplotigs’, was done with the purge_haplotig v.1.1.1 package (Roach, et al. 2018). The assembly graph generated by *hifiasm* was converted to a set of contigs in multi-fasta format using AWK (Aho, et al. 1987) as described at https://github.com/chhylp123/hifiasm. We assessed focal species containment by using BlobTools2 (Challis, et al. 2020) to detect assembly contamination by non-focal species. In order to quantify biological completeness of our contig set, we used the package BUSCO v.4.1.4 (Manni, et al. 2021) with the eudicotyledons_odb10 ancestral lineage dataset.

### Scaffolding with Hi-C and Bionano

To scaffold contigs generated by *hifiasm*, we used a paired approach of Bionano optical map construction and Hi-C scaffolding. Research has suggested that higher accuracy scaffolding can be attained by first applying Bionano hybrid scaffolding (Bickhart, et al. 2017). A Bionano optical map as well as the construction of hybrid scaffolds was made using the Bionano Access software package. We used HiC-Pro v.3.1.0 (Servant, et al. 2015) and the HiTC v1.42.0 R package (Servant, et al. 2012) to visualize the Hi-C contact map for the resulting Bionano scaffolds. Inspection of this contact map identified Bionano scaffolds that could be joined into chromosome-level scaffolds (Figure S1), as well as one misassembly (see above). Although the Hi-C data provided a clear signal for connecting Bionano scaffolds, it did not provide compelling evidence for how those scaffolds should be oriented within chromosomes. Therefore, in this final assembly step, which connected 11 Bionano scaffolds into the remaining four chromosomes, we determined order and orientation based on the expectations that telomeric repeat sequences should be placed at chromosome ends and that annotated gene density should be higher at the chromosome ends than internal centromeric regions. The final Hi-C based scaffolding was performed with a custom Perl script. For Bionano scaffolding, gap sizes were estimated based on the optical map and include inferred locations of the interspersed 6-bp Bionano nicking sites. For the seven scaffold connections inferred from Hi-C data, we used an arbitrary gap size of 100 bp.

### Genome annotation

For annotation, we relied on a combination of protein and transcript evidence constructed from PacBio Iso-Seq and bulk Illumina RNA-seq for our focal species as well as related species in the tribe *Sileneae*. Specifically, we used the Iso-Seq transcriptome data for *S. conica, S. latifolia, S. noctiflora, S. vulgaris*, and *Agrostemma githago* described in Williams, et al. (2021) and Warren, et al. (2023) and Illumina RNA-seq from Havird et al. (2017). Because we relied on multiple annotation approaches, the way these datasets were incorporated differed slightly. To generate protein evidence as input for the different annotation approaches, we used TransDecoder v.5.5.0 to identify the most likely protein-coding regions for individual transcripts in each of the Iso-Seq datasets. To reduce redundancy in our total protein hint dataset, we combined proteins for each individual species and ran CD-HIT v.4.8.1 (Li and Godzik 2006). The resulting protein fasta file was used as the protein hint dataset for all annotation approaches.

Our genome was first soft-masked using a *de novo* generated repeat library created with RepeatModeler2 (Flynn et al. 2020). Only instances of known transposable elements were masked in order to avoid the false masking of genic regions. Transposable element and interspersed repeat content was summarized for visualization with EDTA v2.0.1 (Ou, et al. 2019), using the --anno 1 and --sensitive 1 options, though these annotations were not used as part of the gene annotation process described below.

We provided Funannotate (Palmer and Stajich 2020) with both Illumina RNA-seq and PacBio Iso-Seq data as part of the *train* function, which utilizes a combination of Trinity (Grabherr et al. 2011) and PASA (Haas et al. 2003) to assemble high-quality transcripts. Next, we used the *predict* function, which utilizes the transcripts generated as part of the *train* function plus the protein evidence described above to parameterize the *ab initio* gene prediction software AUGUSTUS (Stanke et al. 2008), which, combined with the alignment of transcript evidence, is then used by Evidence Modeler (Haas et al. 2008) to generate high-quality, consensus gene models.

In order to identify gene models with the BRAKER (Hoff et al. 2019) pipeline we followed the tutorial described at https://github.com/Gaius-Augustus/BRAKER/blob/master/docs/long_reads/long_read_protocol.md. Specifically, after aligning our Illumina short-read data to our genome with STAR (Dobin et al. 2013), we used the resulting BAM file as an input to BRAKER1 which then uses a combination of GeneMark-ES/ET/EP (Bruna et al. 2020) and AUGUSTUS to generate gene models. Next, we used our protein evidence to generate a second set of gene models using BRAKER2. Finally, following the collapsing of redundant transcripts in our PacBio Iso-Seq data for *S. conica* using cDNA_Cupcake (https://github.com/Magdoll/cDNA_Cupcake), we used TSEBRA (Gabriel et al. 2021) to both compare the gene models generated with BRAKER1 and BRAKER2 to our Iso-Seq based gene models and also retain the best amongst the three sets of evidence.

Finally, we used the full set of transcripts for *S. conica* and the optimized AUGUSTUS models generated as part of the Funannotate pipeline and the protein evidence described before, as inputs for the MAKER2 pipeline (Holt and Yandell 2011). The full set of configuration files used for four separate iterations of MAKER2 are available at on Github (https://github.com/Sloan-Lab/Silene_conica_genome_project). We included a separate mapping iteration, two iterations with a combination of AUGUSTUS and SNAP (Johnson et al. 2008), and an iteration with GeneMark-ES/ET/EP (Bruna et al. 2020). The resultant gene models were filtered to retain those which had an AED score ≤ 1.

We used the software AGAT (Dainat et al. 2023), specifically the function *agat_sp_merge_annotations*.*pl*, in order to merge and de-duplicate annotations generated by the three separate approaches. We used the AGAT function *agat_sp_keep_longest_isoform*.*pl* to remove isoforms from our annotation.

### Analysis of DNA transfer between nuclear and cytoplasmic genomes

To identify sequences transferred between the nuclear and cytoplasmic genomes, we searched published *S. conica* mitochondrial and plastid genome sequences (Sloan, Alverson, Chuckalovcak, et al. 2012; Sloan, Alverson, Wu, et al. 2012) against our nuclear genome assembly, using NCBI BLASTN v2.12.0+ with the *-task blastn* option. BLAST hit locations and the percentage of the nuclear and cytoplasmic genomes that were covered by hits (e-value threshold of 1e-6) were summarized with custom Perl scripts (https://github.com/Sloan-Lab/Silene_conica_genome_project). A sliding window analysis was also performed to summarize the percentage of shared sequence in 1-Mb windows (with a 250-kb step size) along the length of the nuclear chromosomes. Coverage values and individual BLAST hits were visualized with a custom R script (https://github.com/Sloan-Lab/Silene_conica_genome_project). Only hits with a minimum length of 300 bp were visualized with individual tick marks, but all hits meeting the e-value threshold (1e-6) were used for calculating and visualizing coverage percentages. These calculations were performed after removing hits from two very large regions of ribosomal DNA repeats (one on Chr6 and the other on Chr8) that result from ancient similarity between nuclear, mitochondrial, and plastid rRNA genes (rather than recent transfers between genomes).

### Methylation analysis

PacBio HiFi sequencing data can also be used to detect some types of epigenetic modifications. We used the ccsmeth v.0.3.2 (Ni, et al. 2022) package to detect 5-methylcytosine base modifications in a CpG context (5mCpGs). Specifically, putative per-base modification information was first detected using the PacBio software CCS v.6.4.0 (flag ‘--hifi-kinetics’; https://github.com/PacificBiosciences/ccs) followed by alignment to the target genome using pbmm2 v1.7.0 (https://github.com/PacificBiosciences/pbmm2). ccsmeth then applies a deep-learning-based model (here, *model_ccsmeth_5mCpG_aggregate_attbigru_b11*.*v2*.*ckpt*) to infer methylation state across the target genome. Analyses with ccsmeth were done using a NVIDIA RTX 3090 graphical processing unit (GPU).

## Data availability

The raw data (PacBio HiFi reads, Illumina-based Hi-C reads, and Bionano optical map data), genome assembly, and annotation can be accessed under the NCBI BioProject PRJNA904366 (assembly version 2: JAQQAY000000000.2). Annotation data can be found in the Zenodo repository at the following DOI: 10.5281/zenodo.8223290. All scripts for bioinformatic analyses are available at https://github.com/Sloan-Lab/Silene_conica_genome_project.

## Acknowledgements

This project was supported bv the National Science Foundation (MCB-2048407) and the National Institutes of Health (R35 GM148134).

---

**Table S1.**
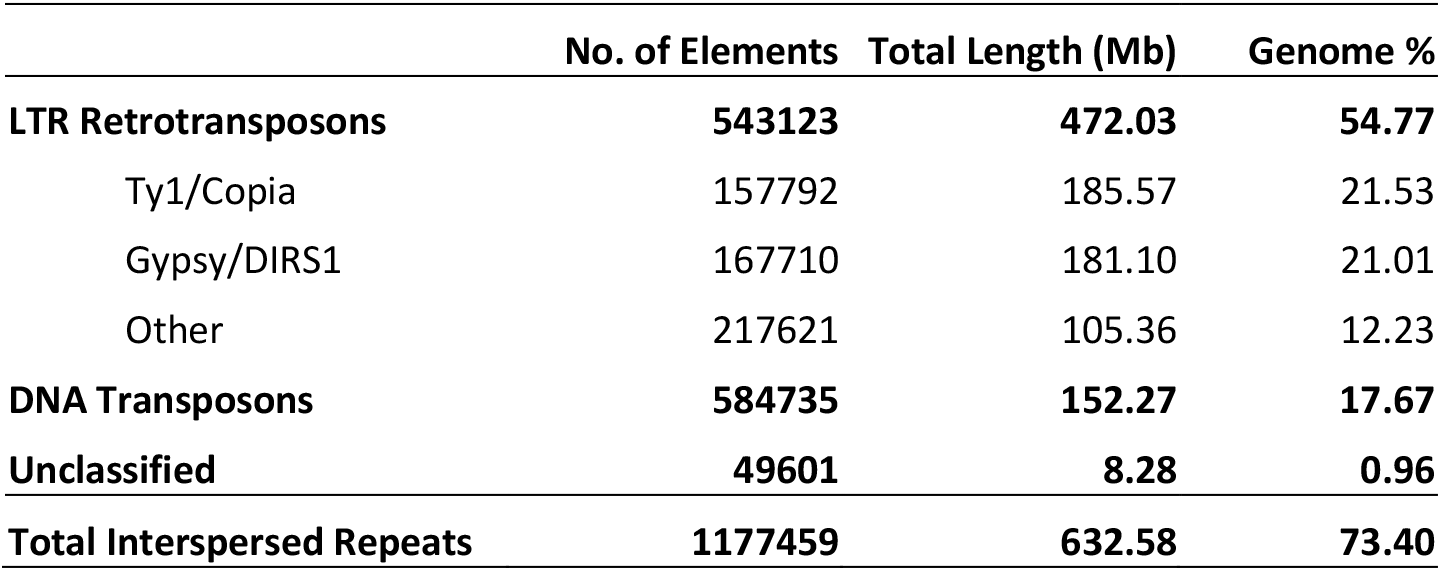
Interspersed repeat content in the *S. conica* genome.

**Figure S1.**
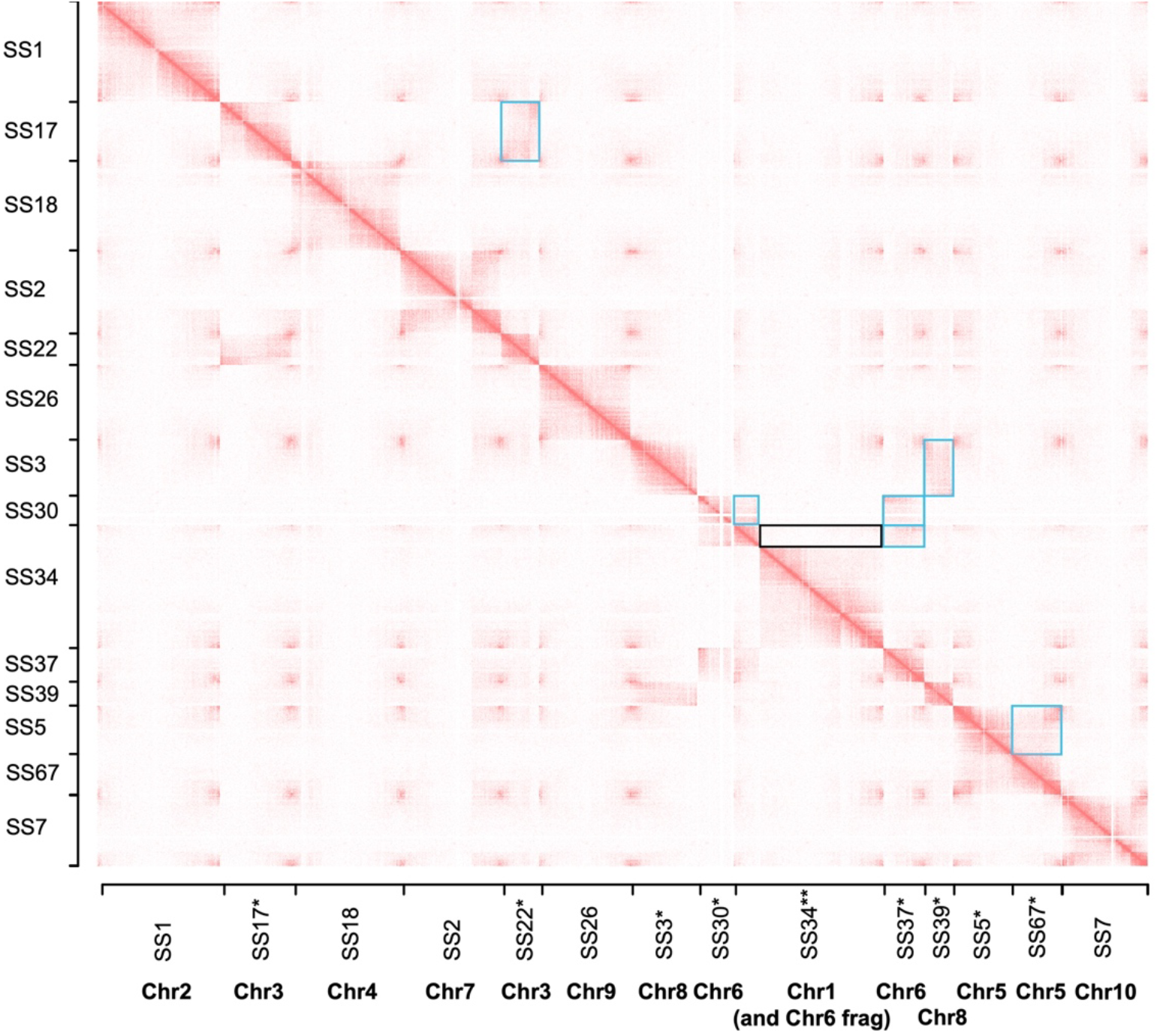
Heatmap generated with HiC-Pro and the HitC R package visualizing Hi-C interactions across the Bionano super-scaffolds (SS) from the *S. conica* genome. The chromosomes to which the scaffolds were eventually assigned are indicated on the x-axis. The outlined boxes above the primary diagonal highlight signal that led to joining scaffolds (teal boxes) or breaking apart a misassembly due to lack of Hi-C contacts within the scaffold (black box). The corresponding regions below the primary diagonal are left unhighlighted for visual comparison. Single asterisks (*) indicate Bionano scaffolds that were joined to form larger chromosome-level scaffolds. The double asterisk (**) indicates the misassembled Bionano SS34 scaffold that was subsequently separated into Chr1 and a portion of Chr6. Only 14 of the 16 Bionano scaffolds are visualized in this figure because the other two were too small and consisted of tandemly repeated ribosomal DNA regions. Those two scaffolds were joined with the other Chr6 scaffolds based on strength of Hi-C contact signal.

**Figure S2.**
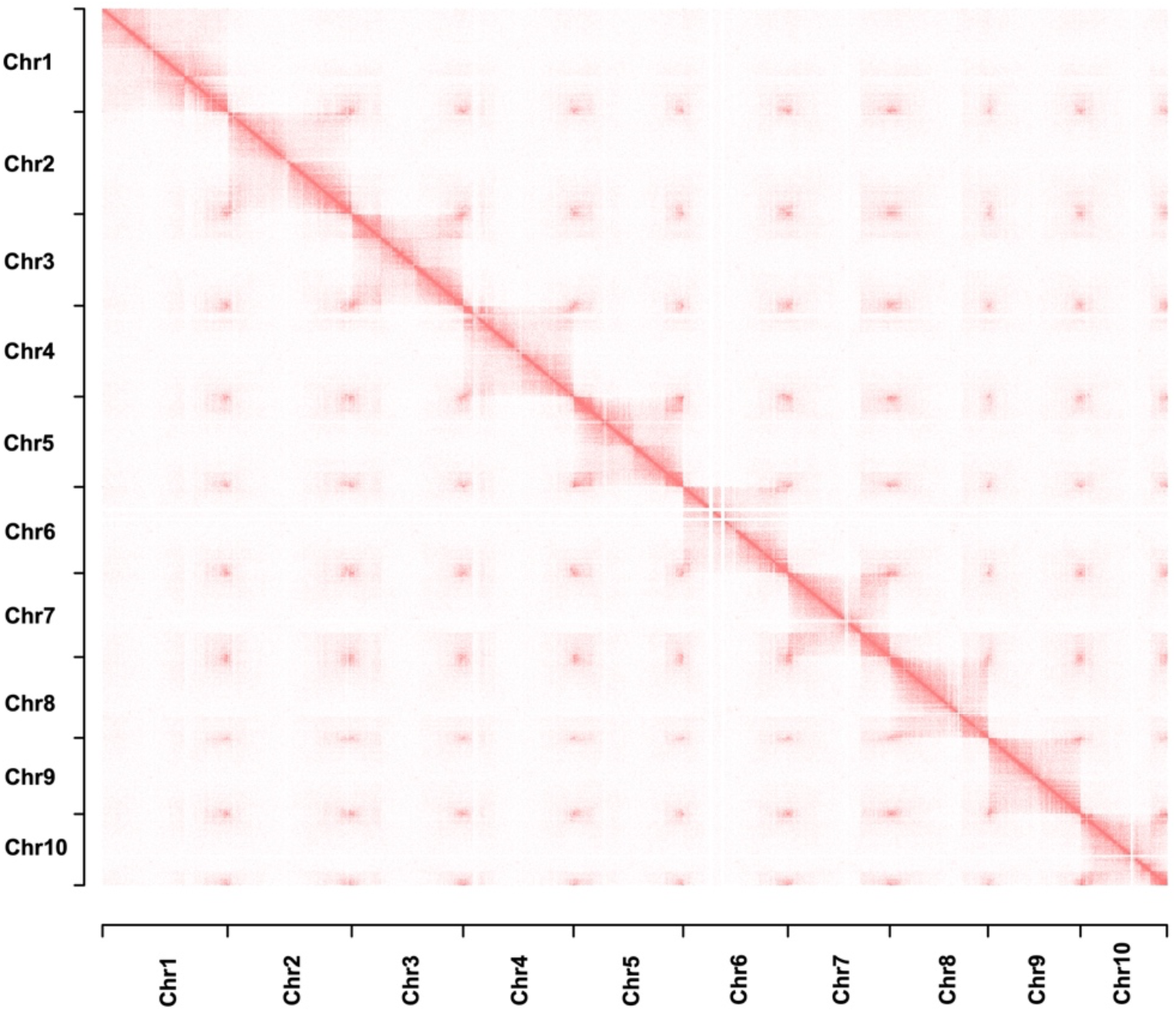
Heatmap generated with HiC-Pro and the HitC R package visualizing Hi-C interactions across the 10 chromosome-level scaffolds from the *S. conica* genome.

